# Lingering representations of stimulus history at encoding influence recall organization

**DOI:** 10.1101/068353

**Authors:** Stephanie C.Y. Chan, Marissa C. Applegate, Neal W Morton, Sean M. Polyn, Kenneth A. Norman

## Abstract

Several prominent theories posit that information about recent experiences lingers in the brain and organizes memories for current experiences, by forming a temporal context that is linked to those memories at encoding. According to these theories, if the thoughts preceding an experience X resemble the thoughts preceding an experience Y, then X and Y should show an elevated probability of being recalled together. We tested this prediction by using multi-voxel pattern analysis (MVPA) of fMRI data to measure neural evidence for lingering processing of preceding stimuli. As predicted, memories encoded with similar lingering thoughts (about the category of preceding stimuli) were more likely to be recalled together, thereby showing that the “fading embers” of previous stimuli help to organize recall.

## Introduction

We have an immense number of memories stored in our brains. Why do we retrieve certain memories at certain times? How are memories organized in the brain and how does this affect recall? These questions have been studied using memory tests such as free recall, in which participants recall items in whatever order they choose. Existing research has uncovered two main organizational phenomena: *semantic contiguity effects* (a tendency for items with similar meanings to be recalled together; Bousfield and Sedgewick 1944; Jenkins and Russell 1952; Romney et al. 1993) and *temporal contiguity effects* (a tendency for items studied close in time to be recalled together; Kahana 1996; Kahana et al. 2008).

Semantic contiguity effects can be explained in terms of participants using features of a just-recalled item as a cue for recalling other items (e.g., if you recall a fruit, you can use retrieved fruit features as a cue to recall another fruit). Temporal contiguity effects require a more complex explanation. Modern *temporal context* theories (e.g., Howard and Kahana 2002) posit that temporal contiguity arises because, at encoding, item representations are linked to a slowly changing context representation. When an item is recalled, it retrieves the context representation that it was linked to at study, which in turn cues retrieval of items that were studied in similar contextual states. Because (by hypothesis) context changes slowly over time, retrieved context preferentially cues items that were studied *close in time* to the just-retrieved item, thus giving rise to temporal contiguity effects. Several studies have found neural evidence for temporal context drift and its influence on memory (e.g., Howard et al. 2012; Hyman et al. 2012; Manning et al. 2012; Polyn et al. 2012).

Some theories are agnostic about what information is contained in this context representation and what causes it to drift (e.g., Estes 1955 and Mensink and Raaijmakers 1988 both posit random drift). However, more recently, theories like the Temporal Context Model (TCM; Howard and Kahana 2002) and the Context Maintenance and Retrieval model (CMR; Polyn et al. 2009) have set forth a more specific account. According to this account, context is composed (at least in part) of lingering information about recently studied items, which are linked to the memory representation of the currently studied item. For example, if you switch from talking about football to politics, then these theories posit that the “fading embers” of your football thoughts will persist in your mind for some time and become linked to your memory for the politics discussion. This view converges with recent neuroscientific data showing that information is represented at multiple time scales in the brain, such that some areas only represent the current focus of attention, whereas other areas integrate over longer time scales (Hasson et al. 2008); it also converges with neurophysiological data on “time cells”, showing that different populations of neurons are involved in representing a stimulus, as a function of how long ago the stimulus was presented (e.g., MacDonald et al. 2011; Howard and Eichenbaum 2013). In essence, models like TCM and CMR posit that neural populations that represent *preceding* stimulus information get linked to the neural populations that represent *current* stimulus information, thereby contextualizing that information.

The signature prediction of this theory is that, if the lingering thoughts active during an experience X are similar to the lingering thoughts active during an experience Y, then the memories of X and Y should show an elevated probability of being recalled together, because they will have been linked to similar (lingering) information. Remarkably, despite this prediction’s centrality, it has not yet (to our knowledge) been tested. In this experiment, we sought to test this prediction by using multi-voxel pattern analysis (MVPA) of fMRI data (Lewis-Peacock and Norman 2014; Norman et al. 2006) to track evidence, at the time of an item’s encoding, for neural representation of the preceding item’s category.

We collected two datasets (n=17 and n=24), following the same data collection procedures for both.^1^ Using a multi-voxel classifier of fMRI data that was designed to pick up on lingering traces of the preceding category, we show that activity patterns reflecting properties of preceding stimuli influenced the organization of recalls, as predicted by theories of temporal context like TCM – that is, memories encoded with similar “lingering thoughts” about the category of preceding items were more likely to later be recalled together during recall.

## Materials and Methods

### Participants

For the first dataset, we recruited 17 participants (aged 18-33 years, 11 female) from the Princeton University community. For the second dataset, we recruited 24 participants (aged 18-29 years, 18 female). All participants provided informed written consent. The study was approved by the Princeton University Institutional Review Board.

### Task

While undergoing functional magnetic resonance imaging (fMRI), participants studied lists of items from different categories. After studying each list, they performed a recall-by-category task, where participants were cued to recall items from specified categories, one category at a time. The lists of items were organized with a particular category structure that allowed us to test how lingering thoughts about preceding items at study affected recall organization at test.

An overview of the task is shown in Fig 1. At the start of each study-test block, participants were presented with a list of 18 items, one at a time. The items belonged to one of three categories: we schematically refer to them as **A**, **B**, and **M** (where M stands for “main”, because these were the main items of interest; the A- and B-items served to contextualize the M-items, as described below). In any given list, the roles of A, B, and M were mapped one-to-one onto the following three categories of pictures: celebrities, landmarks, and objects. For example, in one list, the A-items might be landmarks, the B-items might be objects, and the M-items might be celebrities. The assignment of categories (celebrity, landmark, object) to roles (A, B, M) was counterbalanced such that, across lists, each category served equally often in the A, B, and M roles.

**Fig 1.**
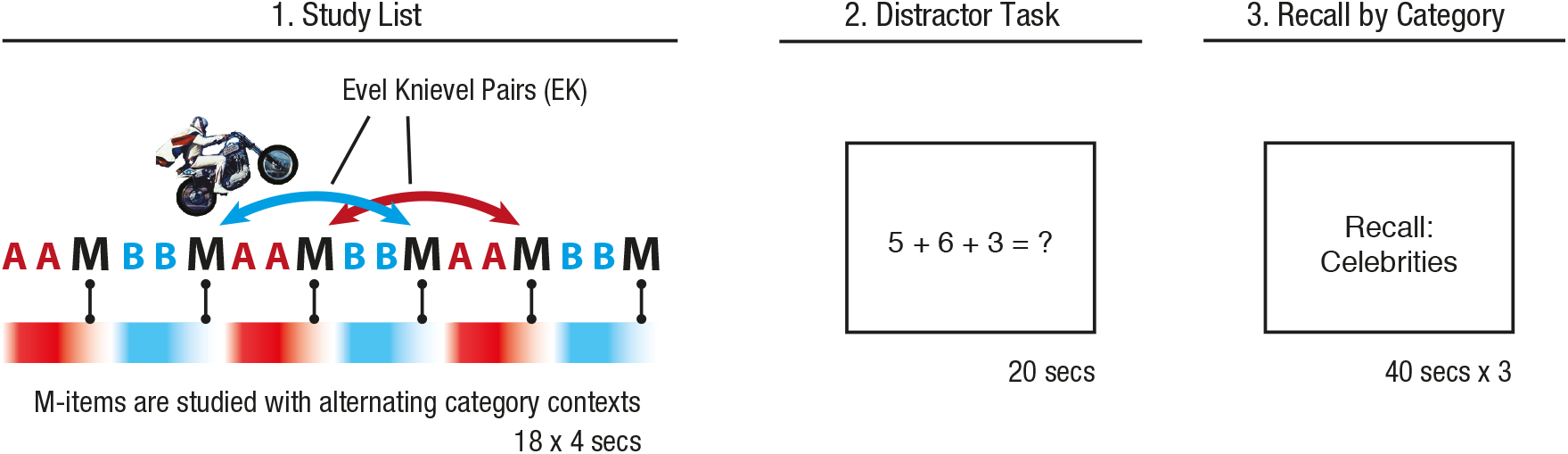
Trial structure for the recall-by-category task. Each trial begins with a study list. 18 items were shown one at a time, every 4 seconds. Each study list was composed of items from three different categories (labeled A, B, and M), and the lists were structured as shown. After the study list, participants performed 20 seconds of a distractor task, followed by recall of the items in the M-category (in this example: celebrities), followed next by recall of items in the A and B categories. “Evel Knievel (EK)” transitions refer to transitions at recall between M-items that were studied with the same preceding category, because they “jump over” temporally nearer M-items (EK transitions may be of length 2 or 4, and may be backwards or forwards).

During the study lists, a new item appeared every 4 seconds, coinciding with the onset of an fMRI image acquisition (each item was shown for 3400 ms, with 600 ms of fixation after each item). Each item presentation was composed of a photograph of a celebrity face, a famous landmark, or a common object, with the name of the item (e.g., “Eiffel Tower”) presented below the photograph; the stimuli were adapted from those used in Morton et al. (2013). To encourage encoding of the items, participants were required to make a category-specific judgment of each item on a 4-point scale (celebrities: “How much do you love or hate this person?”; landmarks: “How much would you like to visit this place?”; objects: “How often do you come across this object in your daily life?” Polyn et al. 2005).

After the presentation of the 18 list items, participants performed 20 seconds of a distractor task (self-paced arithmetic problems – summing three random digits, multiple-choice with four choices).

After the distractor task, participants were asked to verbally recall as many items from the list as possible, one category at a time; within each category, participants were allowed to recall freely (i.e., in any order). Participants were first asked to recall M-items (“main items”), and then the A- and B-items; participants were given 40 seconds to recall each category. We analyzed recall data only from the M-items, but we asked participants to recall the A- and B-items as well, to ensure that they paid attention to those items during study.

There were 12 study-test blocks in total. The experiment task was coded using Psychtoolbox 3 (http://psychtoolbox.org). The verbal recalls for the M-items were annotated using Penn TotalRecall (http://memory.psych.upenn.edu/TotalRecall).

The primary dependent measure of interest was the order in which M-items were recalled, as manifested in patterns of *recall transitions*. We say a “transition” has occurred from item X to item Y when participants recall items X and then Y in immediate succession (without recalling any intervening items).

The key to our study-list structure was that the M-items were preceded by *context items* that alternated in category (A then B then A then B…) (see Fig 1). According to temporal context theories like TCM, the M-items should be linked to lingering thoughts about the preceding category (either A or B), and this linking to the preceding category should influence the organization of recall. In the absence of this influence, temporal contiguity effects should dominate the patterns of recall, favoring recall transitions between neighboring M-items, as has been previously observed for free recall experiments using study-lists without this alternating semantic structure (e.g., Kahana 1996; Polyn et al. 2011). However, if lingering category information is indeed “contextualizing” M-items in our study-lists, there should be a boost in transition probability between M-items that were preceded by the same category. Because the A and B context items alternated in category, these transitions between M-items with matching preceding-category context involve “leaping over” a temporally nearer M-item in favor of a farther M-item; accordingly, we call these transitions “Evel Knievels” (or *EK transitions*), after the daredevil stuntman famous for his motorcycle jumps across canyons, piled cars, and other obstacles. EK transitions could be of length 2 or 4 (jumping over 1 or 3 M-items), and in the forward or backward directions.

### Overview of fMRI analysis

As noted above, our main hypothesis was that lingering thoughts relating to preceding items would become linked to M-items at study, thereby resulting in an elevated probability of transitions between M-items that were preceded by the same “context” category (i.e., EK transitions). Importantly, we also expected there to be moment-to-moment variability in the extent to which preceding-category information was represented in participants’ brains; we only expected to see a boost in EK transitions for the subset of trials where preceding-category information actually persisted in participants’ brains. To test this prediction, we used fMRI pattern classifiers (Lewis-Peacock and Norman 2014; Rissman and Wagner 2012) to track participants’ thoughts about the preceding category. By estimating the level of lingering category information associated with particular M-items, we could make predictions about the order in which these M-items would later be recalled. Specifically, we predicted that — for a pair of M-items X and Y that were preceded by the same context category and could thus be later recalled together as an EK transition — preceding-category information for X and Y (as measured by the classifier) would be more similar when participants actually made the EK transition, compared to when they made a non-EK transition instead.

As an additional analysis, we used the same logic to address whether the properties of the M-items themselves affected recall order. Previous work suggests that participants might also use information about the semantic category of the items themselves (in addition to retrieved context information) to cue memory recall (manifesting as semantic contiguity effects) (Rissman and Wagner 2012; Morton et al. 2013). If so, items associated with strong category-specific activity during encoding would be more likely to be clustered together during recall. To measure this potential second effect on recall organization by the semantics of the items themselves (as opposed to the preceding items), we used fMRI pattern classifiers to also measure the amount of M-category information elicited by each M-item; we refer to this as current-category information, to distinguish it from preceding-category information. Following the same logic as our main analysis, we investigated whether—for the same pair of M-items X and Y that could later be recalled together as an EK transition—levels of current-category match for X and Y were higher when participants actually made the EK transition vs. when they did not.

### fMRI acquisition and pre-processing

Functional brain images were acquired using a 3T MRI scanner (Siemens, Skyra) and were preprocessed using FSL (http://fsl.fmrib.ox.ac.uk/fsl/). An echoplanar imaging sequence was used to acquire 40 slices (3mm iso, repetition time (TR) = 2s, echo time (TE) = 30ms, flip angle = 71º). We collected 3 study-test blocks in each scanning run; there were 4 scanning runs in total. The functional images were spatially filtered using a Gaussian kernel (full width at half maximum of 5mm) and temporally filtered using a high-pass cutoff of 0.0077Hz. We performed motion correction using a six-parameter rigid body transformation to co-register functional scans, and then registered the functional scans to an anatomical scan using a 6-parameter affine transformation. Data were spatially normalized by warping each participant’s anatomical image to MNI space using a 12-parameter affine transformation. To prepare the data for pattern classification, the activity for each voxel was z-scored within each study-test block.

### MVPA classifier training and testing

Multi-voxel pattern analysis (MVPA) was performed using the Princeton MVPA Toolbox (https://code.google.com/p/princeton-mvpa-toolbox/). We trained two distinct pattern classifiers. First, we trained a classifier to decode information about the category of the current stimulus. Second, we trained a classifier to decode lingering information about the category of the preceding stimuli, based on neural activity from the time of the current stimulus. We trained two distinct classifiers (instead of using just one classifier to decode both current and preceding stimulus identity) because of recent evidence (mentioned above: Hasson et al. 2008; Howard and Eichenbaum 2013) suggesting that different neural populations may be responsible for coding the current stimulus vs. lingering information about preceding stimuli. The training methods for these two distinct classifier types are described below.

To create training and testing examples for the classifier designed to detect the category of the *current* stimulus, we labeled each brain image with the category of the stimulus presented at that time. Because brain images were acquired every 2 seconds and stimuli were presented every 4 seconds, each stimulus was linked to two brain images. Then we shifted these labels 4 seconds forward in time; this shift accounts for lag in the hemodynamic response measured by fMRI. For example, if the participant studied a celebrity for 4 seconds, then the two images acquired starting 4 seconds and 6 seconds after the onset of the celebrity were labeled as being “celebrity” brain patterns (see Fig 2, top).

**Fig 2.**
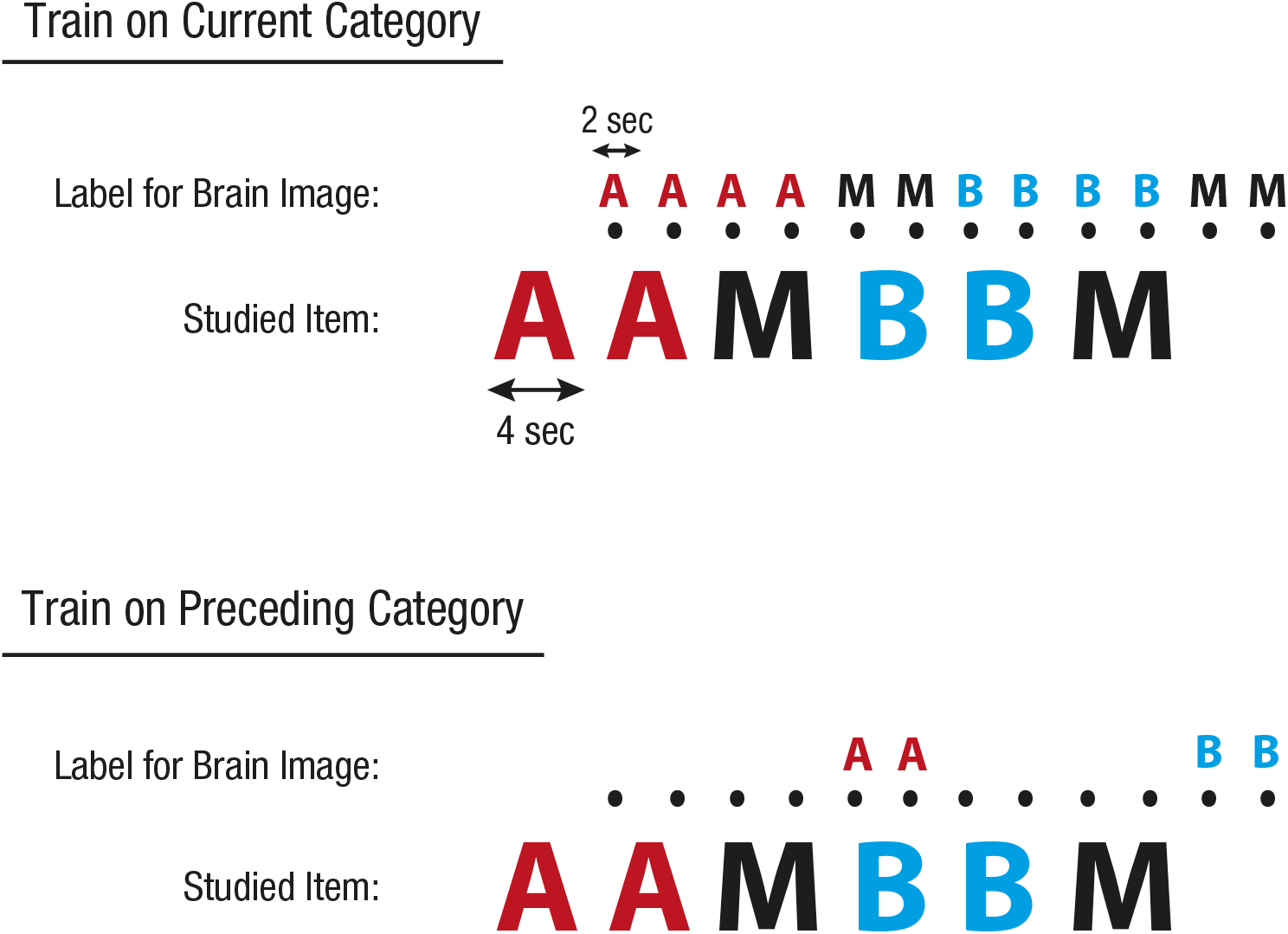
Labeling of brain images for MVPA classifiers. We trained and tested MVPA classifiers in two different ways: (1) training and testing on the current semantic category; (2) training and testing on the preceding semantic category. The figure illustrates how brain images (indicated by dots) were labeled for the two classifier types. Brain images were collected every two seconds; stimuli were presented every four seconds (stimulus onset was timed to coincide with the start of an image acquisition). See text for additional differences between our preceding-category classifiers and standard current-category classifiers.

To create training and testing examples for the classifier designed to detect the category of the *preceding* stimulus, we took the brain images for which we would expect the peak response to each M-item (the same brain images that we used to train a classifier on the *current* category, acquired 4 and 6 seconds after the onset of the M-item), and — instead of labeling those images with the category of the M-item (as we did above) — we labeled those images with the category that *preceded* that M-item. For example, if the M-item was a celebrity that was preceded by landmarks, we would label those images as being “landmark” brain patterns (see Fig 2, bottom). All other (unlabeled) images were left out of classifier training and testing.

For each participant, we trained three separate preceding-category classifiers – one classifier for the lists where the M-category was celebrities, one for the lists where the M-category was landmarks, and one for the lists where the M-category was objects. In this way, the classifiers could not use current-category information to aid in classifying the preceding category, since the current category was held constant for all training (and testing) examples. To further aid the classifier in focusing on preceding-category information, we used feature selection that selected against voxels that varied significantly with the current category (ANOVA-based feature selection with a threshold of *p* = 0.05). The next section describes in more detail the rationale for the design of the preceding-category classifier.

For both current-category and preceding-category classifiers, we used logistic regression with L2 regularization (using a regularization penalty of 1; classifier performance was not very sensitive to this parameter). Specifically, we trained a logistic regression classifier for each category to respond with a “1” when an image was labeled with that category and with a “0” when an image was not labeled with that category. Once trained and presented with new input data, these category-specific classifiers output a real value from zero to 1, indicating the degree of neural evidence for the category that it was trained to detect. Classifiers were always trained and tested in a leave-one-block-out fashion — e.g., to apply the classifier to a time point from study-test block 1, the classifier was trained on blocks 2 through 12.

### Validating the preceding-category classifier

Our initial procedure for training a preceding-category classifier (originally applied to the first dataset) produced classifiers that in fact opportunistically used current-category information to aid in that classification, so that the classifier outputs reflected information that we did not intend to incorporate. Here, we describe the corrected procedure we used to create an improved preceding-category classifier, and we show why it is superior to the more straightforward approach that we originally used.

When training preceding-category classifiers, we initially did not take any measures to hold the current category constant across lists – we trained a single preceding-category classifier for all lists, rather than training three separate classifiers for M-as-celebrity lists, M-as-landmark lists, and M-as-object lists. As a result, classifiers trained on preceding-category labels could in fact opportunistically use current-category information to aid in the classification of the preceding-category. When current category is not controlled in the set of training images, information about the current category informs the classifier about what the preceding category is *not*. Figure 3a shows the output of the current-category classifier, averaged over all lists. The negative weighting against the current category is visible when we examine timecourses of output from our original preceding-category classifier (Fig 3b). It is especially apparent in the first few timepoints of the study list – these timepoints are not preceded by any A, B, or M items, and so we should expect a true preceding-category classifier to be at chance. However, the classifier has learned that the current category cannot be the same as the preceding-category, and consequently shows a clear negative bias against A.

**Fig 3.**
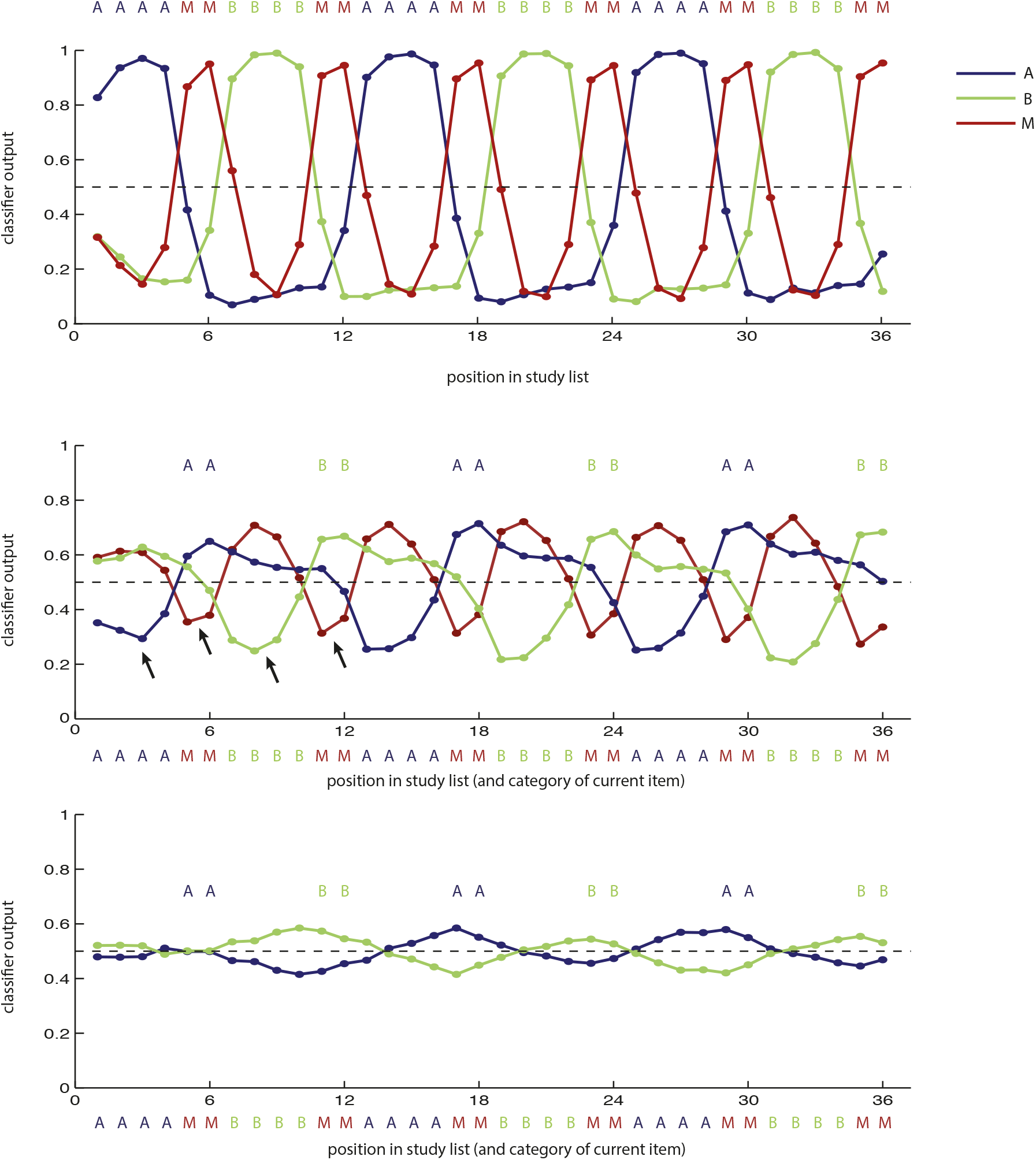
Timecourses of output for the MVPA logistic regression classifiers, averaged across lists and participants. Colored letters above each plot indicate the training labels. Colored letters below each plot indicate the category of the current stimulus (after correcting for hemodynamic lag). (a) Outputs for classifiers trained to identify the current category. (b) Outputs for classifiers that were trained on M-timepoints from all but one list to identify the preceding category, and then applied to all timepoints of the remaining list. This version was trained on celebrity, landmark, and object lists together. These classifiers show bias against the current-category (black arrows indicate a few examples of negative activation of the current category). (c) Same as b, except this classifier was trained and tested separately for lists with the M-category as celebrity, landmarks, and objects (so that the current category was held constant for the training examples for each classifier), used a whole-brain mask instead of a temporal-occipital mask, and used feature selection against voxels with strong current-category information. These classifiers no longer show a consistent bias against the current-category. They also show a gradual buildup of the A and B categories that peaks at the first TR of each M-item.

To remedy this problem, we made three changes to the classifier. Firstly, we trained three separate classifiers for each participant: one classifier for the lists where the M-category was celebrities, one where the M-category was landmarks, and one where the M-category was objects. In this way, information about the current category was not available to the classifier, since the current-category was held constant for all training (and testing) examples for each classifier (remember that we only used the M-timepoints for preceding-category classification). Secondly, we used a whole-brain mask instead of a temporal-occipital mask, to allow the classifier to draw from more anterior parts of the brain, in case persistent information about recent stimuli is represented there (previous research has shown that this does appear to be the case, e.g. Hasson et al. 2008). Lastly, we implemented feature selection that selected *against* voxels that varied significantly with the current category (we removed these voxels from consideration, using ANOVA-based feature selection with a threshold of *p* = 0.05).

This classifier training procedure is disadvantaged in that it only has 3 lists for each cross-validated training iteration (rather than 11), and may suffer from having less data. However, as can be seen in Fig 3c, this new version of the classifier has a very different profile from the one in Fig 3b, and no longer shows the same bias against the current category. In fact, as we would expect, these classifiers generally show outputs that slowly ramp up through each block of A- or B-items, peaking at the 1^st^ TR for each M-item.

### Using classifier evidence to compute current-category and preceding-category match for pairs of items

We predicted that, if two M-items were studied with similar profiles of preceding category information, participants would be more likely to transition directly between these items at recall (this directly tests the hypothesis that preceding-category information contextualizes the M-items in memory).

To evaluate this prediction, we computed the *preceding-category match* (PCM) for each pair of M-items. PCM measures how much a given pair of M-items registered as being preceded by the same category context.

Preceding-category match (PCM) for pair of M-items was computed as:

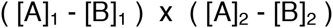

where [A]_1_ is the level of A-category evidence at the time of studying the 1st M-item in the pair, [A]_2_ is the level of A-category evidence at the time of studying the 2nd M-item in the pair, and so on. Importantly, A-category and B-category evidence in this score was read out using classifiers trained to detect the *preceding* category, described above. The subtractions [A]_1_ - [B]_1_ and [A]_2_ - [B]_2_ measure the “balance” of lingering category evidence (in favor of A vs. B) for the 1st and 2nd M-items. If both M-items strongly favor the same preceding category (both favor A or both favor B, i.e. [A] - [B] for both M-items is strongly positive or strongly negative), then the PCM score is strongly positive (close to +1). In such cases, we would expect a relatively high probability of recall transition between the two M-items, because the MVPA decoders indicate that the M-items were encoded with matching preceding-category contexts. If the M-items strongly favor opposite categories (one favors A and one favors B), then the PCM score is strongly negative (close to -1). In such cases, we would expect a relatively low probability of recall transition between the two M-items (Fig 4a).

**Fig 4.**
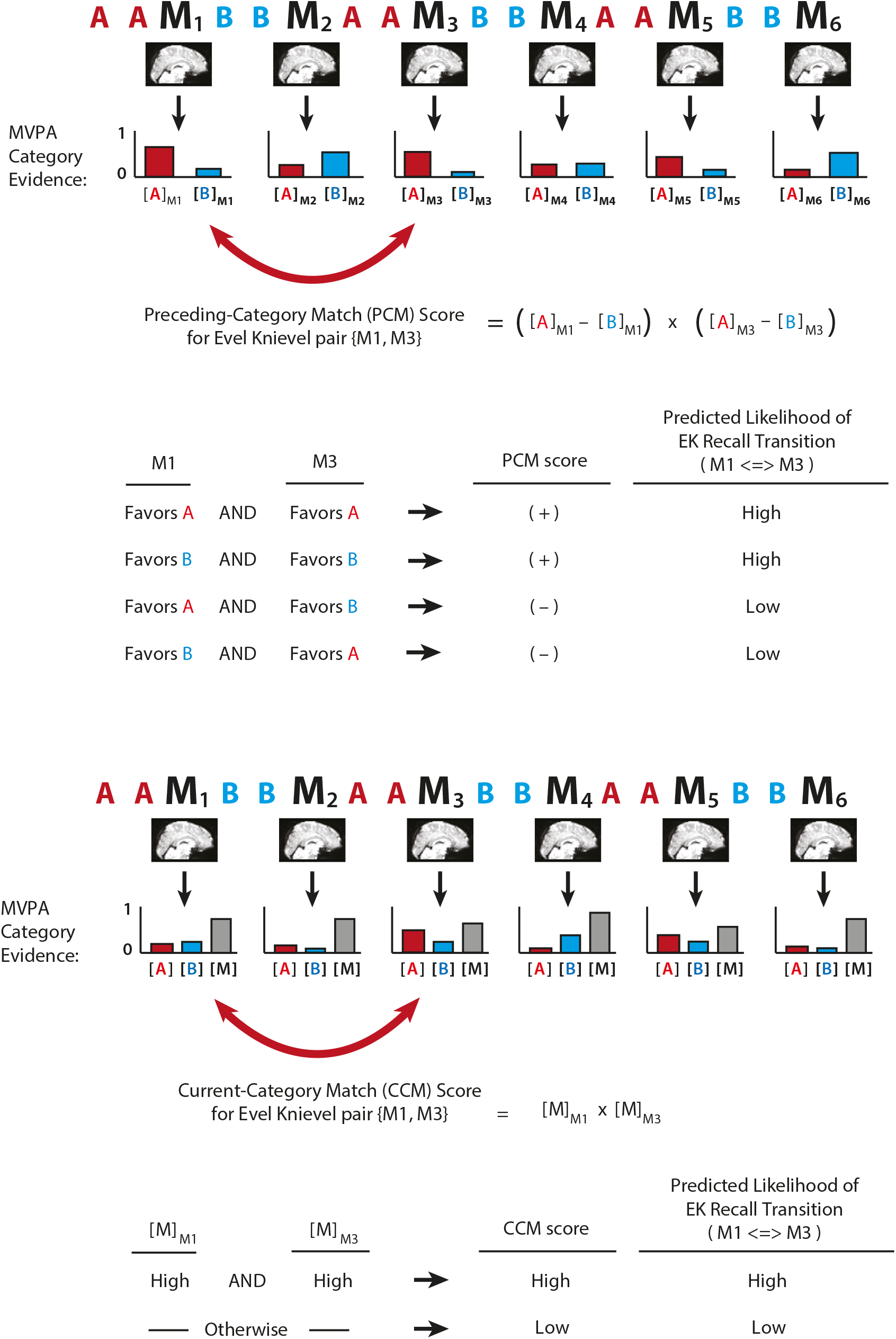
Procedure for computing preceding-category and current-category match scores. (a) MVPA analysis to compute preceding-category match (PCM) for a pair of M-items. Classifiers were trained to identify the preceding category. Classifier outputs were interpreted as levels of evidence for each category. For a given pair of M-items, outputs from these classifiers were combined to form the PCM score, which was designed to measure the degree to which preceding-category information matched for the two M-items. (b) MVPA analysis to compute current-category match (CCM) for potential EK transitions. Classifiers were trained to identify the current category. For a given pair of M-items, these classifier outputs were multiplied to obtain the CCM score, which was designed to measure the degree to which both M-items triggered neural activity corresponding the current category.

We also evaluated the degree to which participants were more likely to recall items together if those items both triggered neural activity corresponding to the *current* category (i.e., basic semantic clustering). Current-category match (CCM) for a pair of M-items was computed as:

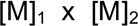

where [M]_1_is the level of M-category evidence associated with the 1st item in the pair, and [M]_2_is the level of M-category evidence associated with the 2nd item in the pair. Importantly, M-category evidence in this score was read out using classifiers trained to detect the *current* category. Previous work showed that items associated with strong category-specific brain activity during encoding tend to be recalled as part of a cluster of same-category items during free recall (Morton et al. 2013). This finding suggests that items associated with strong category-specific activity provide good retrieval cues for one another. Therefore, we also investigated whether a given pair of items would be more likely to be recalled together if their CCM score was high (Fig 4b), indicating strong category-specific activity during encoding of both items.

### Relating classifier evidence to recall order

To test our predictions about how recall order depends on match in preceding-category representations, we looked at recall of M-items, and separated the observed recall transitions into EK and non-EK transitions. Our goal was to assess whether there were reliable differences in preceding-category match (PCM) for potential EK pairs when participants “jumped over” a nearer M-item to make the EK transition, vs. when they made a non-EK transition to the just-nearer M-item. We predicted that, when comparing PCM for a potential EK pair vs. PCM for the just-nearer nonEK pair, this difference would be higher when participants actually made the EK transition during recall (vs. when they instead made the non-EK transition).

We also performed a parallel analysis using current-category match (CCM) instead of preceding-category match (PCM), to evaluate any effects of current-category match on recall organization.

To ensure that we carried out a fair comparison between EK and non-EK transitions (Fig 5), we only analyzed EK transitions where it was actually possible for participants to instead have made a non-EK transition in the same direction, to the just-nearer M-item (i.e., the just-nearer M-item had not already been recalled). Likewise, we only analyzed non-EK transitions where it was actually possible for participants to instead have made an EK transition in the same direction, to the just-farther M-item. We excluded non-EK transitions where the just-farther M-item had already been recalled, and we also excluded non-EK transitions where participants transitioned backward to the first M-item or forward to the last M-item on the list (in these cases, there *was* no just-farther M-item). Because of this extra exclusion condition for non-EK transitions, we ended up excluding more non-EK transitions than EK transitions: on average, we excluded 17% of EK transitions (95% CI: 13-22%) and 46% of non-EK transitions (95% CI: 41-51%).

**Fig 5.**
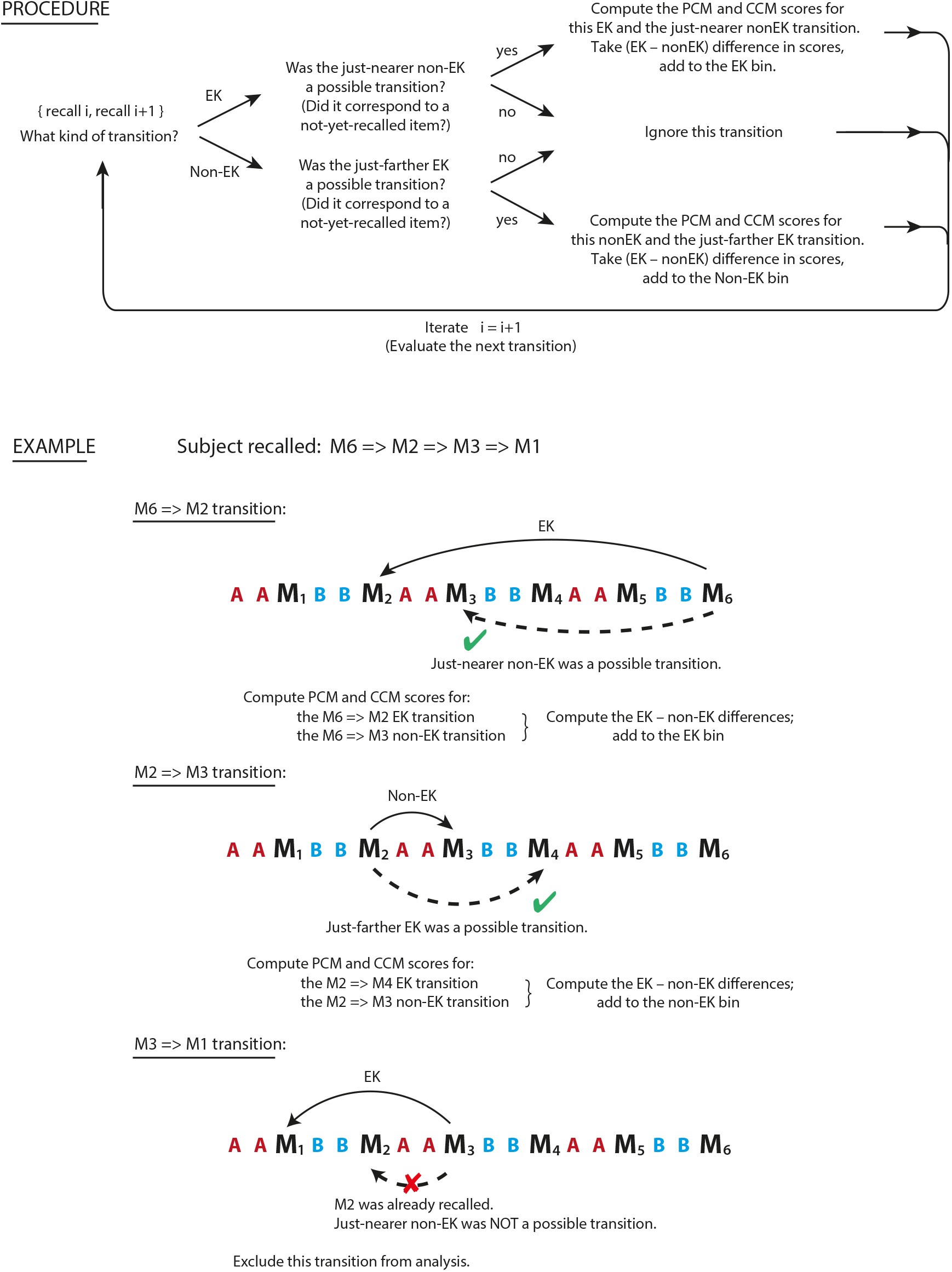
Procedure for aggregating valid EK and non-EK transitions, to ensure fair comparison between the two transition types. In our main analysis, we only included EK transitions where a non-EK transition to the just-nearer M-item would also have been possible (i.e., the just-nearer M-item had not already been recalled), and we only included non-EK transitions where an EK transition to the just-farther M-item would also have been possible. If transitions to the just-nearer M-item (for an EK transition) or the just-farther M-item (for a non-EK transition) were not possible, then we ignored this transition and continued to the next. Otherwise, we considered it a valid transition and included it in our analysis. For EK transitions, we computed the match scores for the EK pair and the just-nearer non-EK pair, and computed the difference. For non-EK transitions, we computed the match scores for the just-farther EK pair and the non-EK pair, and computed the difference. The figure shows an example recall sequence (M6, M2, M3, M1) for a particular list; for this sequence, we would include M6=>M2 as a valid EK transition, include M2=>M3 as a valid non-EK transition, and exclude M3=>M1 as an invalid EK transition.

In order to capture the relative strength of preceding- (or current-) category match for a potential EK pair, compared to its just-nearer potential non-EK pair, we computed PCM (or CCM) for both pairs of items and took the difference between the two scores.

### Statistics and confidence intervals

For all of our analyses looking (separately) at behavioral data or neural data, we computed random-effects bootstrap confidence intervals on the mean by resampling participants with replacement (Efron and Tibshirani 1986). When assessing differences between conditions, we computed bootstrap confidence intervals on the difference between the means. In the text, these are reported as 95% confidence intervals. In the results figures, these bootstrap distributions and confidence intervals are displayed using cat’s eye plots.

## Results

### Behavioral results

On average, participants correctly recalled 54.8% of the M-items that they studied (95% CI: 51.5–58.1%). Broken down by category, participants recalled 63.4% of celebrity M-items (95% CI: 60.0–66.8%), 60.5% of landmark M-items (95% CI: 55.6% to–65.2%), and 40.4% of object M-items (95% CI: 36.1–44.6%). Participants complied with our instructions not to repeat themselves during free recall (i.e., they never recalled the same M-item twice during a single recall period). Participants occasionally made intrusions (i.e., recalled items not on the current study list); transitions involving intrusions were not included in our EK analysis (e.g., if a participant recalled item M2, an intrusion next, and finally item M4 after that, then neither the M2=>intrusion nor the intrusion=>M4 transitions were included in our analysis). On average, each participant made 0.13 intrusions per list (95% CI: 0.092–0.18). Of these intrusions, 26% on average were the names of items studied on previous lists (95% CI: 14–42%); the other intrusions were names that had not appeared anywhere in the experiment. On average, each subject made 10.4 valid EK transitions (95% CI: 9.0–11.8) and 16.6 valid non-EK transitions (95% CI: 15.0–18.6), where “valid” is as defined above and in Fig 5. These behavioral results are reported in Table 1 for each dataset individually.

**Table 1.** Behavioral results for each dataset individually (95% confidence intervals in parentheses).

|  | Dataset 1 (n=17) | Dataset 2 (n=24) |
| --- | --- | --- |
| % of M-items correctly recalled | 55.7% (51.9 - 59.6%) | 54.2% (49.2 - 59.2%) |
| % of celebrity M-items correctly recalled | 64.2% (59.6 - 69.1%) | 62.8% (58.0 - 67.5%) |
| % of landmark M-items correctly recalled | 62.0% (55.8 - 68.0%) | 59.4% (52.6 - 66.3%) |
| % of object M-items correctly recalled | 40.9% (36.57 - 45.3%) | 40.0% (33.4 - 46.6%) |
| mean number of intrusions per list | 0.132 (0.078 - 0.196) | 0.132 (0.083 - 0.212) |
| % of intrusions that were prior-list items | 29.6% (11.1 - 66.7%) | 23.7% (10.5 - 47.4%) |
| mean # of valid EK transitions | 10.6 (9.0 - 12.6) | 10.3 (8.5 - 12.1) |
| mean # of valid non-EK transitions | 17.1 (15.1 - 18.9) | 16.3 (14.0 - 19.4) |

### Basic classifier results

Before relating the classifier output to recall behavior, we first wanted to establish that the preceding-category classifier was decoding category identities at above-chance levels.

For the classifiers trained to decode the preceding category, we computed accuracy for each fMRI image based on whether classifier evidence for the correct context category (A or B: whichever one actually preceded this particular M-item) was greater than classifier evidence for the incorrect context category. For this 2-way classification, chance is 50%. The observed level of accuracy was 57% for lists with celebrities as the M-category (95% CI: 54-60%), 57% for lists with landmarks as the M-category (95% CI: 53-61%), and 58% for lists with objects as the M-category (95% CI: 54-61%). (These classifier results are reported in Table 2 for the individual datasets.) Importantly, these accuracy percentages only denote the percentage of outputs that matched the preceding-category *labels* that we provided to the classifier—not the match to the participants’ actual neural activity. We believe that the output of the classifier in fact reflects a noisy estimate of meaningful fluctuations in the extent to which preceding-category information lingered in participants’ brains. In our main analysis, this variability in the classifier output is what allows us to make predictions about when participants will make EK transitions.

**Table 2.** Basic preceding-category classifier results for each dataset individually.

| M category | Dataset 1 (n=17) | Dataset 2 (n=24) |
| --- | --- | --- |
| celebrities | 57% (54 - 59%) | 57% (54 - 60%) |
| landmarks | 58% (55 - 62%) | 56% (53 - 59%) |
| objects | 57% (54 - 60%) | 58% (55 - 60%) |

Reported are mean classifier accuracies (and 95% confidence intervals) for the three different preceding-category classifiers.

### Relating classifier evidence to recall order

As noted above (Fig 5), our analysis of recall order focused on recalls where participants had the opportunity to make either an EK transition or a just-nearer non-EK transition. In keeping with our predictions, preceding-category match (PCM) scores predicted participants’ recall behavior: The difference in PCM scores for the EK pair vs. the just-nearer non-EK pair (Fig 6a, top) was larger when participants made the EK transition vs. when they made the non-EK transition. That is, participants were more likely to recall two M-items together if the M-items were encoded with matching lingering information about preceding items. This result provides direct support for the idea that lingering thoughts relating to preceding items serve to contextualize memories and organize subsequent recall. Considering the individual datasets: for the second dataset, the result was significant on its own; for the first dataset, the pattern was qualitatively similar, but the effect size was smaller overall and the effect did not reach significance (Fig 6a, middle and bottom).

**Fig 6.**
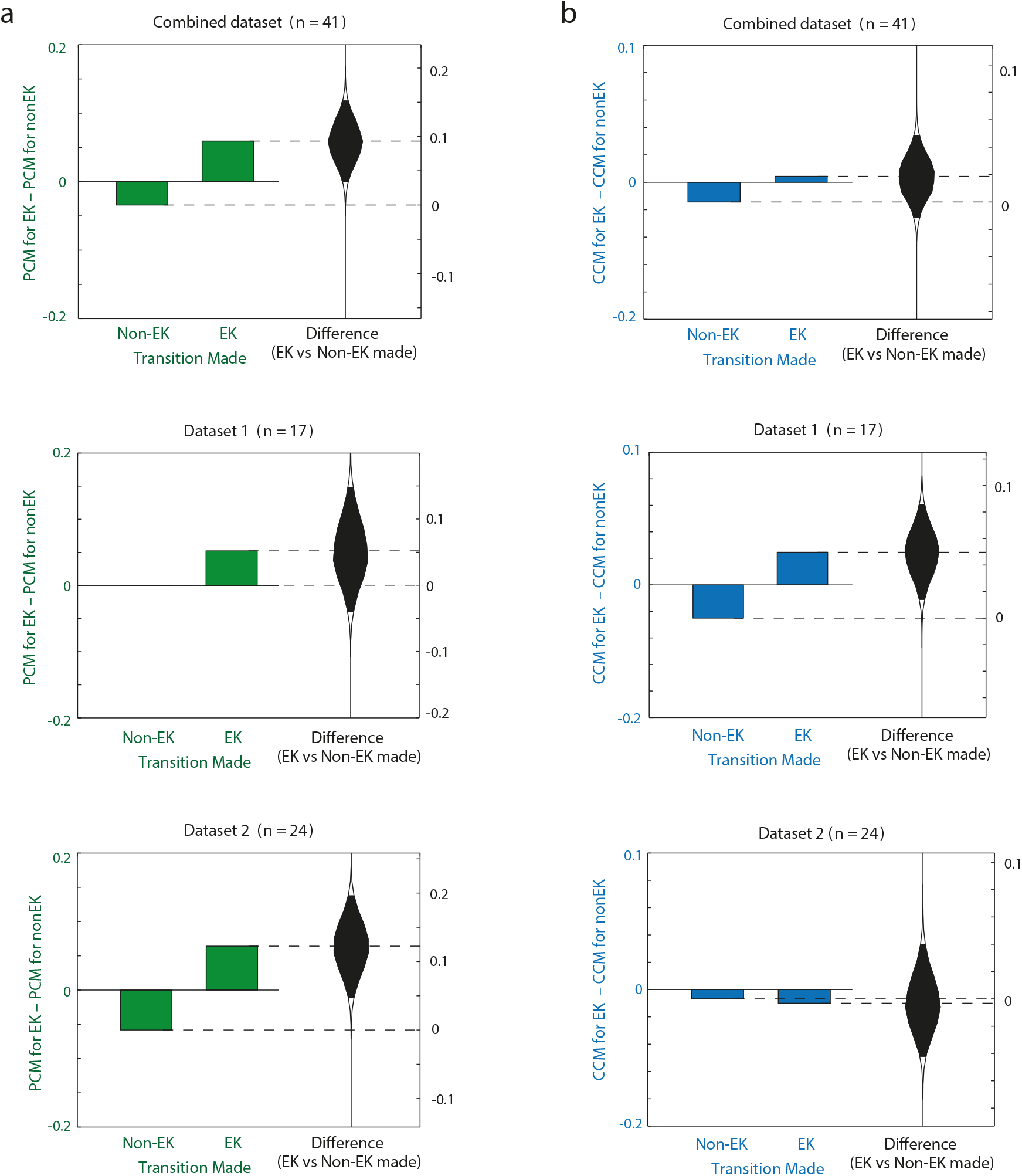
Main results. Bars indicate (a) the difference in preceding-category match (PCM) scores for EK and just-nearer non-EK pairs, (b) the difference in current-category match (CCM) scores for EK and just-nearer non-EK pairs, as a function of whether participants ended up making the EK transition or the non-EK transition. Cat’s-eye plots show bootstrap distributions for the difference in the bars. Results are shown for the combined data, and for the individual datasets. Black shaded areas of cat’s-eye plots indicate 95% confidence intervals. For the combined dataset, PCM scores significantly predicted recall behavior but CCM scores did not.

We did not find a corresponding effect of current-category match on recall order in the combined dataset (Fig 6b, top). When analyzing individual datasets, we did find a current-category effect in the first dataset, but not in the second dataset (Fig 6b, middle and bottom). The findings from the second dataset, where there was a substantial, significant PCM effect and a near-zero CCM effect (numerically in the opposite direction) indicate that – at least for that dataset – the preceding-category results were not driven by current-category match.

Table 3 reports effect sizes for how well preceding-category match and current-category match predicted EK transitions (these effects correspond to the cat’s eye plots in Fig 6).

**Table 3.**
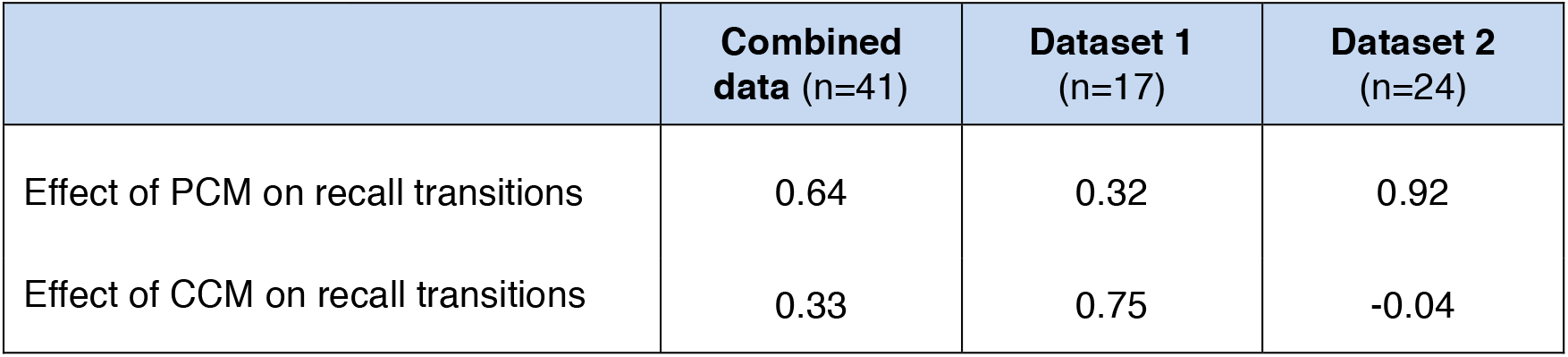
Effect sizes (reported as Cohen’s d) for effect of preceding-category match (PCM) scores and current-category match (CCM) scores on recall transitions.

## Discussion

In this study, we used fMRI pattern classification to track lingering traces of preceding thoughts, and we showed that memories encoded with similar “lingering thoughts” about the category of preceding items were more likely to later be recalled together. The idea that items are contextualized by the “fading embers” of recently studied items is a central assertion of extant models of temporal context and memory (e.g., Howard and Kahana 2002). Our results provide the most direct evidence to date in support of this view.

These effects of “lingering thoughts” on recall order are distinct from previously documented effects of *current* stimulus properties on recall order. For example, Morton et al. (2013) found that the degree of category-specific activity elicited by a studied item predicted category clustering on a free recall test (see also Kuhl et al. 2012, who found that category-specific activity at encoding predicted cued recall success at test). In our combined dataset, we did not find that information about the current item’s category significantly predicted which items would later be recalled successively. In the second study, the size of the effect of preceding category match (measured using Cohen’s d) was 0.92 and the size of the effect of current category match was 0.04 in the wrong direction. This indicates that information about the *preceding* item’s category exerted influences that were not driven by, and were distinct from, any potential influences of the current item’s category. It is possible that the lack of a robust current-category-match effect in our study was due to our use of a *recall-by-category* paradigm, where participants were instructed to recall only one category at a time; in this situation, recalls are always clustered by category, so there is less variance in recall order for the neural measure to explain (relative to free recall, where items can be recalled in any order). In our study, we thought that items that registered neurally as “better” exemplars of a category (according to the current-category classifier) might be more likely to be recalled together, but that effect was not reliably observed here.

Our results, taken together with previous experiments, point to a synthesis whereby multiple time scales of representation influence recall organization, in distinct ways. As noted above, previous studies have shown that information about what is *currently* happening gets encoded into the memory trace, leading to semantic clustering effects (e.g. Morton et al. 2013). The main contribution of the present study is to show that information about what was *recently* happening is also encoded into the memory trace. Under normal circumstances, this can lead to temporal clustering (if participants see events A, B, C in sequence, lingering information about A gets encoded along with both B and C, leading to enhanced transitions between B and C). In our study, however, we deliberately structured study lists so that the effects of lingering semantics on recall could be investigated separately from other potential origins of temporal clustering effects. In our study lists, encoding of preceding-category information worked *against* temporal clustering — to the extent that participants were integrating preceding-category information into their memory traces, they should make “Evel Knievel” recall transitions that jump over nearer items. This is exactly what we saw.

One limitation of our study is that we only tracked thoughts relating to the *immediately preceding* category. As such, our results do not, on their own, discriminate between dual-store memory models (which posit that recently studied items linger in a short-term memory store, so that adjacent items are directly associated with each other during study; Atkinson and Shiffrin 1968; Raaijmakers and Shiffrin 1981) and retrieved-context models like TCM (which posit that items are associated with a separate temporal context representation that contains information about recently presented items; Howard and Kahana 2002; Polyn et al. 2009). It is worth emphasizing that both of these accounts (dual-store models and retrieved-context models) posit that activity relating to preceding items persists in some form and is linked in memory to the current item. Thus, the effect of lingering information on recall organization is a key prediction of a set of prominent theories of memory, for which our experiment is the most direct test to date.

1 The second dataset was originally collected to replicate a result found in the first dataset. After collection of both datasets, we discovered an error in our original analysis, and here use a corrected analysis to analyze both datasets (the original and corrected analyses are described in “Validating the preceding-category classifier” of the Methods). In this paper, in addition to showing the results for the combined data, we also show results for each dataset individually.

